# Antibiotics damage the colonic mucus barrier in a microbiota-independent manner

**DOI:** 10.1101/2024.03.19.585540

**Authors:** Jasmin Sawaed, Lilach Zelik, Yehonatan Levin, Rachel Feeney, Maria Naama, Ateret Gordon, Mor Zigdon, Elad Rubin, Shahar Telpaz, Sonia Modilevsky, Shira Ben-Simon, Aya Awad, Sarina Harshuk-Shabso, Meital Nuriel-Ohayon, Michal Werbner, Bjoern O Schroeder, Amir Erez, Shai Bel

## Abstract

Antibiotic use is a risk factor for development of inflammatory bowel diseases (IBDs). IBDs are characterized by a damaged mucus layer, which does not properly separate the host intestinal epithelium from the microbiota. Here, we hypothesized that antibiotics might affect the integrity of the mucus barrier. By systematically determining the effects of different antibiotics on mucus layer penetrability we found that oral antibiotic treatment led to breakdown of the mucus barrier and penetration of bacteria into the mucus layer. Using fecal microbiota transplant, RNA sequencing followed by machine learning and *ex vivo* mucus secretion measurements, we determined that antibiotic treatment induces ER stress in the colonic tissue which inhibits colonic mucus secretion in a microbiota-independent manner. This mucus secretion flaw led to penetration of bacteria into the colonic mucus layer, translocation of microbial antigens into circulation and exacerbation of ulcerations in a mouse model of IBD. Thus, antibiotic use might predispose to development of intestinal inflammation by impeding mucus production.

## Introduction

Antibiotics are a broad family of drugs that disrupt multiple crucial processes in microbes. Since their discovery, antibiotics have become life-saving therapeutics used to treat microbial infections. The prolific use of antibiotics in both medicine and agriculture has resulted in the rise of antibiotic-resistant microbes, which pose a major challenge to modern healthcare^1,2^. This extensive use of antibiotics is based on the assumption that, other than toxicity issues when used in large doses, antibiotics disrupt biological processes in microbes and not the host. Yet recent research in germ-free animals is beginning to uncover the overlooked effects that antibiotics have on the host^3–6^.

The growing exposure to antibiotics in the past centuries has been linked to multiple diseases which are now common in industrialized countries. For example, an interaction between diet and antibiotic-induced alteration to the gut microbiota is associated with obesity and diabetes^1,2^. Another group of diseases with rising prevalence in the industrialized world are inflammatory bowel diseases (IBDs)^7^. While the exact etiology of IBDs is not clear^8^, recent epidemiological studies have shown a strong and dose-dependent link between these diseases and antibiotic use^9,10^. Indeed, studies in mice have shown that nutritional changes together with antibiotic use can drive intestinal inflammation^11^. Yet the exact mechanism is not completely understood.

The colonic mucus layer separates the host from the trillions of microbes that inhabit the gut lumen^12^. If this mucus barrier is breached, bacteria can encroach on the host intestinal epithelium and trigger a proinflammatory response^13^. Indeed, breakdown of this barrier is a hallmark of IBDs, and perhaps a driving factor in the development of these diseases^14–16^. Antibiotic treatment in mice leads to translocation and uptake of bacteria to gut-draining lymph nodes while predisposing to development of intestinal inflammation^17^. Yet whether antibiotics directly damage the mucus barrier is not clear. Here, we set out to test the hypothesis that antibiotics predispose to development of intestinal inflammation by disrupting the mucus barrier.

## Results

### Oral antibiotic treatment disrupts the colonic mucus barrier

We orally treated mice with antibiotics to determine whether antibiotic treatment affects the mucus barrier. To mimic short-term antibiotic treatment in patients, we treated the mice twice a day for three days. We used 4 different antibiotics, each belonging to a different class of antibiotics: ampicillin (aminopenicillin class), metronidazole (nitroimidazole class), neomycin (aminoglycoside class) and vancomycin (glycopeptide class). To quantify bacterial penetration into the colonic mucus barrier we fixed the tissues in Carnoy’s fixative, which preserves the mucus barrier and the native bacterial spatial localization^18,19^, and stained bacteria using a pan-bacterial fluorescent *in situ* hybridization (FISH) probe. We found that all 4 antibiotics tested led to breakdown of the mucus barrier and encroachment of bacteria upon the colonic epithelium (Figure 1A and B). Spatial fluorescent intensity imaging also revealed bacterial signals originating in the colonic epithelium (Figure 1C). Thus, short-term oral antibiotic treatment leads to disruption of the mucus barrier.

**Figure 1:**
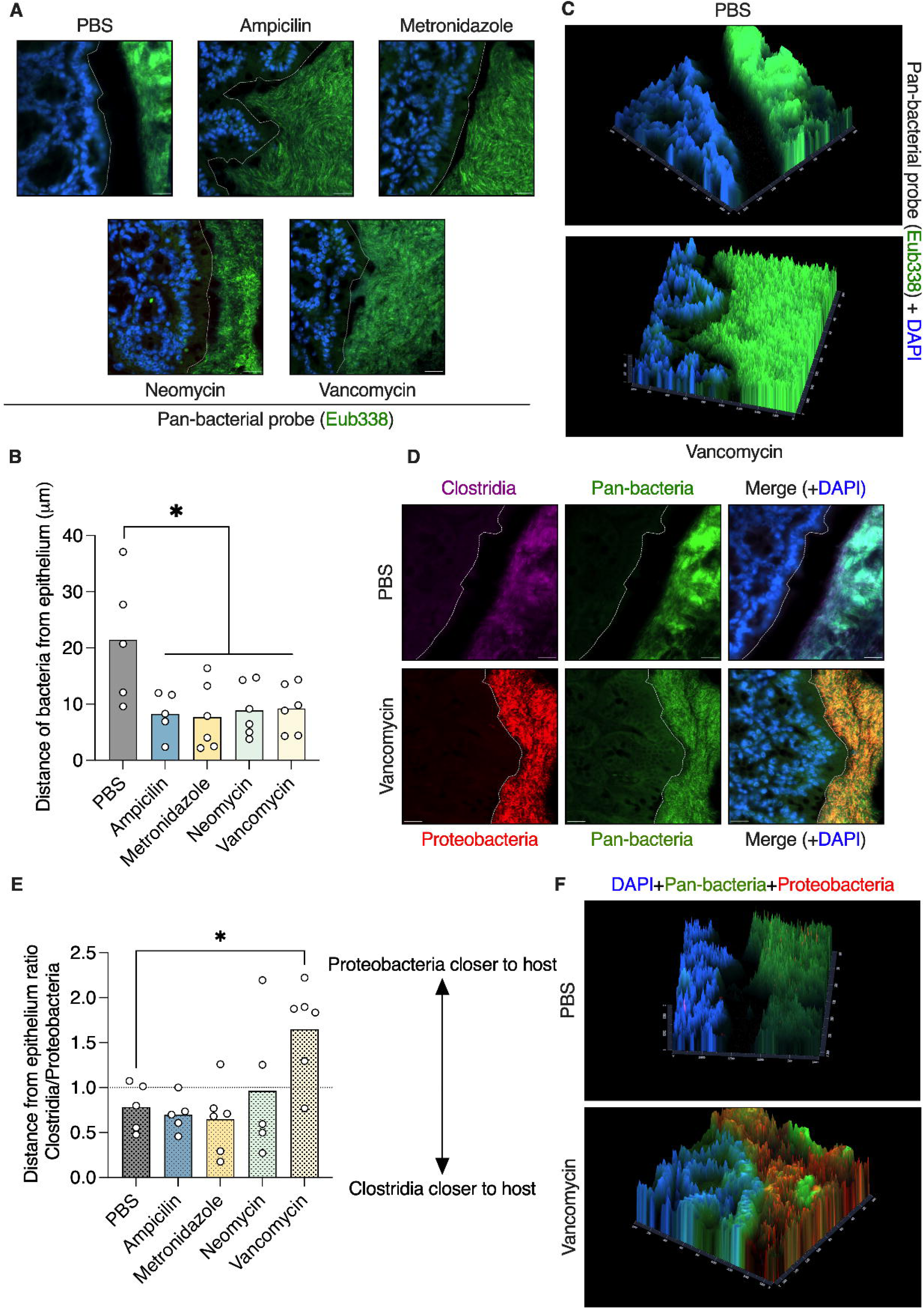
Oral antibiotic treatment disrupts the colonic mucus barrier. (**A**) FISH images of colonic tissues from mice treated orally with antibiotics as indicated. Bacteria are stained in green and host nuclei in blue. The dashed white lines mark the edge of the host epithelium. Scale bars, 20 µm. (**B**) Quantification of distance between luminal bacteria and host epithelium as in **A**. (**C**) Fluorescent intensity imaging of colonic sections from mice treated as indicated. Bacteria are represented by green signal and host epithelium by blue signal. (**D**) FISH images of colonic tissues from mice treated with antibiotics as indicated and stained with the indicated probes. The dashed white lines mark the edge of the host epithelium. Scale bars, 20µm. (**E**) Quantification of the ratio of the distances between Clostridia or Gammaproteobacteria and the host epithelium as in **D**. (**F**) Fluorescent intensity imaging of colonic sections from mice treated as indicated. Pan-bacteria are represented by green signal, Gammaproteobacteria by red signal and host epithelium by blue signal. (**B** and **E**) Each dot represents a mouse. One-way ANOVA. *p<0.05. FISH, fluorescent *in situ* hybridization.

Next, we wanted to identify which bacteria come in close contact with the host epithelium after antibiotic treatment. To this end we performed FISH staining using probes specific to either the Clostridia or the Gammaproteobacteria classes. We chose Clostridia as they are the major Gram-positive class of the gut commensal microbiota, and Gammaproteobacteria as representatives of Gram-negative bacteria which are associated with antibiotic-induced dysbiosis and are associated with multiple diseases^1,2^. In agreement with a previous report^18^ we found that bacteria from the Clostridia class were spatially closer to the host than Gammaproteobacteria in PBS-treated mice (Figure 1D and E). Ampicillin, metronidazole and neomycin treatments did not affect this spatial segregation between Clostridia and Gammaproteobacteria. Vancomycin treatment, which targets Gram-positive bacteria such as Clostridia and is known to cause a bloom of Gammaproteobacteria, led to close contact of Gammaproteobacteria with the host epithelium (Figure 1D-F). Thus, vancomycin is unique as it reverses the spatial position of different bacterial groups in the colon.

### The effect of antibiotics on microbiota composition can not explain antibiotic’s effect on the mucus barrier

Next, we wanted to determine how antibiotic treatment grants the gut microbes access to the niche nearest to the host epithelium. We hypothesized that the effects of antibiotics on microbiota composition disrupts certain microbial communities, thus allowing others to move closer to the host. Because vancomycin is a narrow spectrum antimicrobial, which leads to a bloom of potentially pathogenic Gammaproteobacteria^18^, we chose to focus on it. We treated mice reared in the SPF facility with PBS or vancomycin, as above, and transferred their gut microbiota to germ-free mice via fecal microbiota transfer (FMT). As above, FISH staining in colonic sections from mice treated with vancomycin showed barrier dysfunction and presence of Gammaproteobacteria in close contact with the host epithelium (Figure 2A-C). However, germ-free mice which received a FMT from vancomycin-treated mice did not show a barrier defect or encroachment of Gammaproteobacteria (Figure 2A-C). Interestingly, expression of epithelial-derived antimicrobial transcripts was induced in both vancomycin-treated mice and GF mice that received the FMT from vancomycin-treated mice (Figure S1A and B). This indicates that while the mucus barrier defect can not be explained by changes to the microbiota, the antimicrobial gene signature is induced in response to the vancomycin-induced proteobacteria bloom. Thus, the effect of vancomycin on the gut microbiota can not explain the penetrance of bacteria to the close vicinity of the host epithelium.

**Figure 2:**
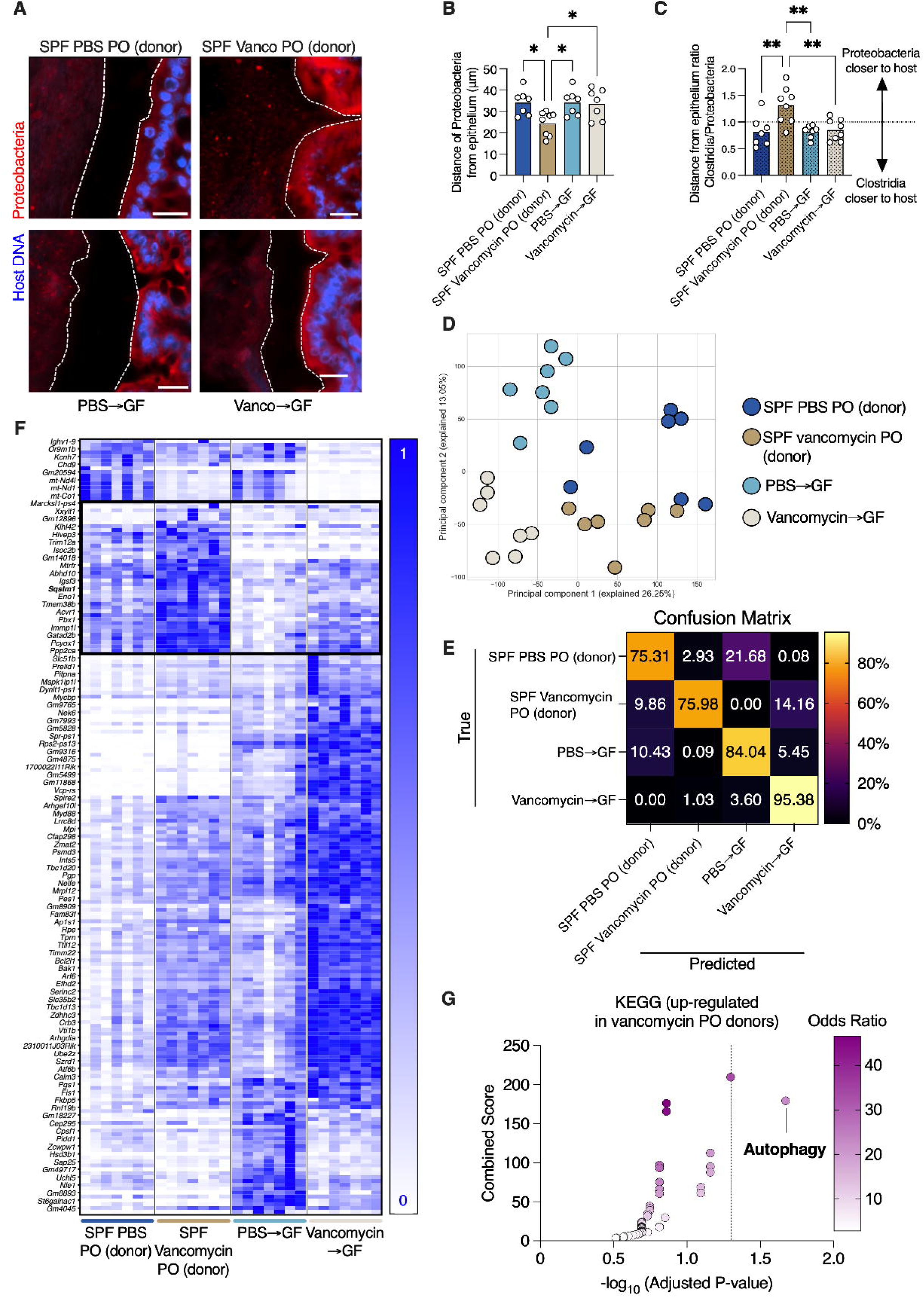
Vancomycin-induced changes to the gut microbiota can not explain treatment impact on the mucus barrier and gut transcription. (**A**) FISH images of colonic tissues from SPF mice treated with vancomycin as indicated or GF mice which received a FMT from vancomycin-treated mice and stained with the indicated probes. The dashed white lines mark the edge of the host epithelium. Scale bars, 20µm. (**B**) Quantification of distance between luminal Gammaproteobacteria and host epithelium as in **A**. (**C**) Quantification of the ratio of the distances between Clostridia or Gammaproteobacteria and the host epithelium as in **A**. (**D**) PCA plot of colonic transcriptional profiles of mice treated as indicated. (**E**) A confusion matrix depicting the percentage of predictions for each category against the true classifications in a 4-way Random Forest classification task. Diagonal entries represent the accuracy of predictions for each category (true positives), while off-diagonal entries indicate the model’s misclassifications. (**F**) Heat map showing transcriptional changes of the top 200 genes which distinguish between the treatment groups based on the classifier in **E**. (**G**) KEGG pathways of transcripts which are uniquely altered in vancomycin-treated mice (donors) as plotted in **F** (in the black box). (**B**-**D**) Each dot represents a mouse. (**B** and **C**) One-way ANOVA. *p<0.05; **p<0.01. FISH, fluorescent *in situ* hybridization; FMT, fecal microbiota transfer; GF, germ-free; PO, *per os*; SPF, specific-pathogen free.

### The effect of antibiotics on host transcription can not be explained solely by antibiotic-induced changes to the gut microbiota

Given our observation that antibiotic-induced changes to the gut microbiota can not explain penetration of bacteria into the mucus barrier, we wanted to test whether vancomycin might be affecting the host directly. To this end, we performed RNA sequencing analysis on colonic tissues from mice raised under SPF conditions that were treated orally with PBS or vancomycin, and germ-free mice that received an FMT from the PBS- or antibiotic-treated mice, as above. In this experimental setup, the SPF mice are exposed to the antibiotic, while the germ-free mice are exposed to the microbial changes which were induced by the antibiotic. We wanted to determine whether vancomycin induced a transcriptional profile in the SPF mice which was unique, compared to the germ-free mice or the SPF mice that received PBS. Therefore, we pursued this clustering using two different approaches. First, we subjected the RNA sequencing data to principal component analysis (PCA). This analysis revealed that each experimental group clustered separately and that the first two principal components accounted for 39% of the variance (Figure 2D). Next, we trained a random-forest (RF) classifier on the 4 groups (SPF vs. GF and PBS vs. Vancomycin) resulting in high confidence classification, as is apparent in the confusion matrix (Figure 2E). The high values along the diagonal of the confusion matrix signify the high percentage of successful assignments of a mouse to the correct group by the RF algorithm. This indicated that each of the four conditions had a unique transcriptional signature, and therefore that there are genes that are activated or suppressed solely in the vancomycin-treated mice. We plotted the gene expression for the 200 most predictive genes according to the RF classifier, with hierarchical clustering separating the different conditions into obvious clusters (Figure 2F). Interestingly, pathway analysis of these genes, selecting those genes that clustered together distinguishing the mice that were orally treated with vancomycin (donors; Figure 2F) using Kyoto Encyclopedia of Genes and Genomes (KEGG) revealed an enrichment in genes related to the autophagy process, including the *Sqstm1* gene which encodes the P62 protein which sequesters cargo for degradation in autophagosomes (Figure 2G). The enhanced expression of autophagy-related genes is consistent with the essential role of autophagy in preserving proper goblet cell function in response to accumulation of ER stress^20^. The fact that autophagy-related genes are activated in response to vancomycin treatment in a microbiota-independent manner, along with the mucus barrier impairment in the mice (Figure 2A-C), implies that vancomycin might be causing stress to goblet cells. Thus, while vancomycin-induced changes to the gut microbiota can explain changes in transcription of many genes, it can not explain all changes to gene expression in mice treated with vancomycin, especially activation of autophagy-related genes.

### Systemic antibiotic administration induces ER stress and disrupts the colonic mucus barrier by inhibiting mucus secretion in a microbiota-independent manner

Given our transcriptomic analysis, which suggested a microbiota-independent effect of vancomycin on the host, we next tested this concept. We hypothesized that antibiotic treatment impairs the mucus barrier via a direct effect on the host. We treated mice with the same four antibiotics as above, but via systemic administration. We found that systemic administration of ampicillin and metronidazole did not significantly change the distance of bacteria from the host epithelium (Figure 3A). This implies that the effects of these two antibiotics on the mucus barrier seen in oral administration (Figure 1A and B) are microbiota-dependent. However, neomycin and vancomycin treatment did impair the mucus barrier (Figure 3A). Interestingly, both neomycin and vancomycin have poor luminal availability when administered systemically, given their inability to cross the gastrointestinal mucosa^21,22^. This further implies that the effect of these antibiotics on the mucus barrier are microbiota-independent. Accordingly, systemic administration of antibiotics did not reverse the spatial position of Clostridia and Gammaproteobacteria (Figure 3B) which was seen with oral treatment above (Figure 1E).

**Figure 3:**
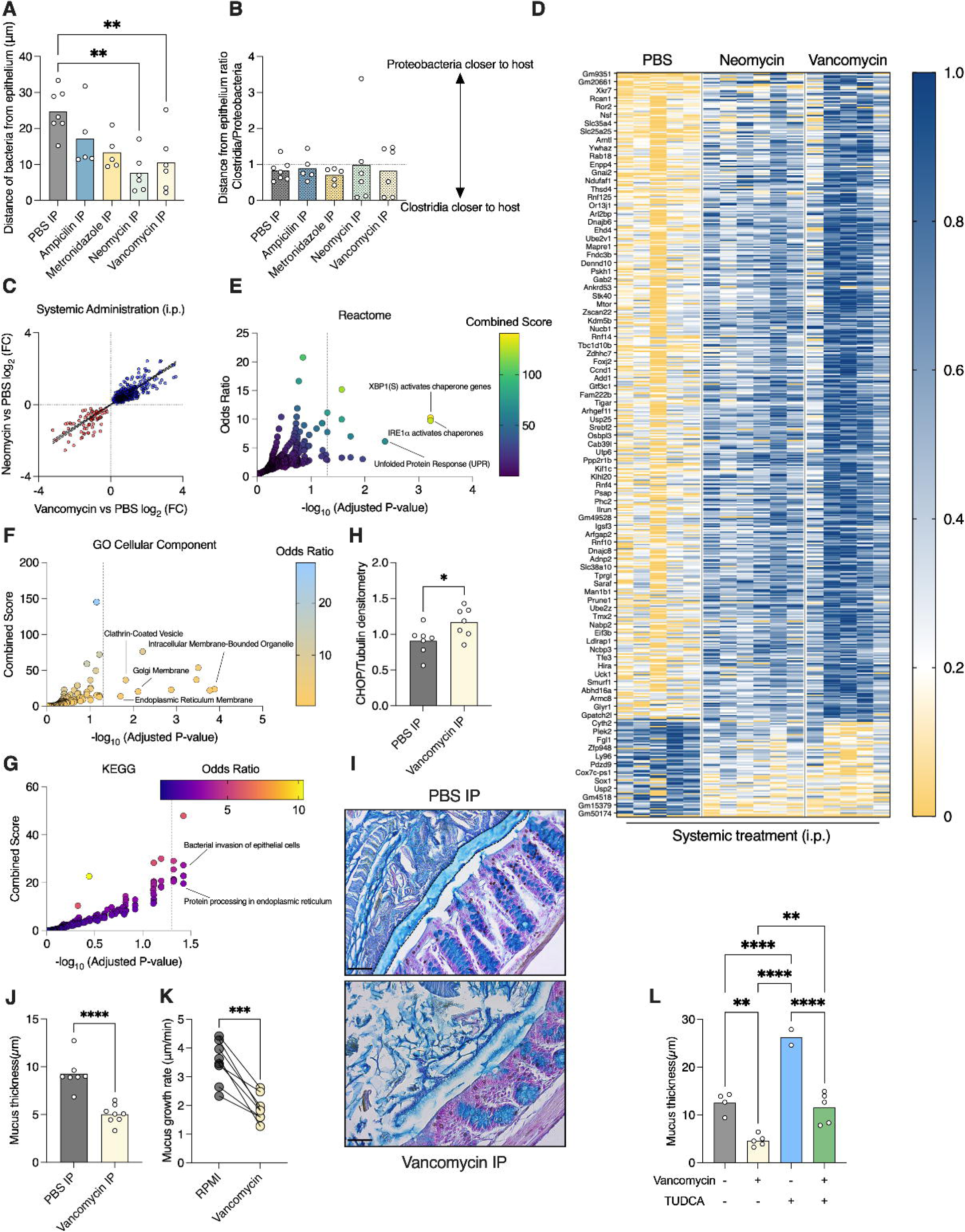
Systemic administration of antibiotics induce ER stress and inhibit mucus secretion in the colon. (**A**) Quantification of distance between luminal bacteria and host epithelium in colons of mice treated with the indicated antibiotics via i.p. One-way ANOVA. (**B**) Quantification of the ratio of the distances between Clostridia or Gammaproteobacteria and the host epithelium in mice treated as in **A**. (**C**) Expression levels of transcripts in colons of mice treated with vancomycin (X axis) or neomycin (Y axis) via i.p. injection as compared to PBS. Plotted are transcripts whose expression was induced by both antibiotics or suppressed by both antibiotics (p<0.05, Mann Whitney test). Each dot represents a transcript. Blue: genes that are more highly expressed under antibiotic treatment; Red: genes with reduced expression after treatment. (**D**) Heatmap depicting normalized expression levels of genes shown in **C**. (**E**-**G**) Analyses using (**E**) Reactome, (**F**) GO cellular component or (**G**) KEGG pathways of transcripts plotted in **C**. Dotted line on the X axis represents a p value of 0.05. Each dot represents a category/pathway. (**H**) Protein levels of CHOP relative to Tubulin using densitometry analysis of a western blot of colonic lysates from mice treated as indicated. (**I**) Colonic sections from mice treated systemically as indicated, stained with Alcian blue to visualize mucus. The mucus layer is defined by the dashed line. Scale bars, 20µm. (**J**) Measurement of mucus thickness as shown in **H**. (**K**) Mucus growth rate. Lines connect tissues from the same mouse. Paired *t* test. (**L**) Measurement of mucus thickness in mice treated as indicated. One-way ANOVA.(**A**, **B**, **H**, **J**, **K** and **L**) Each dot represents a mouse. Student’s *t* test. *p<0.05, **p<0.01, ***p<0.001, ****p<0.0001. IP, intraperitoneal; FC, fold-change; GO, Gene Ontology.

Next, we wanted to determine how systemic neomycin and vancomycin treatment leads to encroachment of the gut microbiota upon the host epithelium. To this end we performed RNA sequencing using colonic tissues from mice treated systemically with either PBS, neomycin or vancomycin. As treatment with these two antibiotics produced the same barrier-dysfunction phenotype (Figure 3A), we tested whether these two antibiotics affect the same transcriptional pathways. We plotted the expression of genes which were induced or suppressed by both antibiotics. To our surprise, we found that the expression of these genes was influenced in a similar manner by the two distinct antibiotics (Figure 3C and D). Pathway analysis revealed that the genes which were similarly affected by the both neomycin and vancomycin are involved in the ER stress response and the unfolded-protein response (UPR) pathways (Figure 3E). Indeed, further analysis predicted that these genes encode proteins which reside in the ER, the Golgi, and secretory vesicles (Figure 3F). Analysis using the KEGG revealed that these genes are activated in response to bacterial invasion to epithelial cells and processing of proteins in the ER (Figure 3G). To verify our RNA sequencing analysis we quantified the levels of the ER stress response protein C/EBP homologous protein (CHOP) and found that it was indeed expressed at higher levels in the colons of mice treated systemically with vancomycin (Figure 3H). Thus, systemic treatment with neomycin or vancomycin induces an ER stress response in the colon.

ER stress is an intracellular switch which limits mucus secretion by goblet cells^20,23^. Given our observation above, that vancomycin induces ER stress in the colon, we hypothesized that vancomycin treatment leads to impairment of the separation between host and microbiota in the colon by inhibiting mucus secretion. We chose to focus on vancomycin as it is used both systemically and orally in the clinic, while neomycin is used mostly topically. To test this hypothesis we treated mice with vancomycin and measured the colonic mucus thickness using Alcian blue staining. We found mice treated systemically with vancomycin lacked a clear mucus in most areas of the colonic epithelial circumference (Figure 3I and J). This observation raised the possibility that vancomycin impairs the ability of goblet cells to secrete mucus. The optimal way to test this hypothesis would be the use of germ-free mice treated with vancomycin. However, germ-free mice do not produce a fully formed mucus layer which can be measured using Alcian blue staining, and also do not contain bacteria to be detected using FISH. To circumvent this problem we measured colonic mucus secretion rates using an *ex vivo* system. We excised colonic sections from naive mice, split them into two sections and fitted them into two measurement chambers. One colonic section from each mouse was infused with media and the other with media supplemented with vancomycin. Using this experimental approach the treated and control tissues originate from the same mouse. We found that all tissues treated with vancomycin showed impaired mucus secretion rates (Figure 3K). As the vancomycin was infused only on the basolateral side of the colonic tissue, and for only 45 minutes, this results demonstrates that the deleterious effect of vancomycin on mucus secretion is microbiota-independent.

Next, we wanted to test whether we could reverse the mucus secretion defect caused by vancomycin treatment. We have previously discovered that the bile acid tauroursodeoxycholic acid (TUDCA) can increase mucus secretion rates by reducing ER stress in colonic goblet cells^20^. As vancomycin treatment induces ER stress in colon (Figure 3E-H) we attempted to restore proper mucus secretion by alleviating this ER stress using TUDCA. Indeed, we found that TUDCA treatment reversed the mucus secretion defect caused by vancomycin treatment, restoring a proper mucus barrier (Figure 3L). Thus, vancomycin treatment inhibits secretion from colonic goblet cells by inducing ER stress.

### Inhibition of mucus secretion by vancomycin use impairs the colonic barrier function and aggravates colonic inflammation

Finally, we wanted to test whether this vancomycin-induced impairment in mucus secretion affects colonic host defense. First, we compared the levels of bacterial antigens in the bloodstream on mice treated systemically with vancomycin. Presence of microbial antigens in the blood is directly linked to intestinal barrier function^24^. We found higher levels of NOD1, NOD2 and TLR5 agonists in the serum of vancomycin-treated mice (Figure 4A-C), indicating impaired colonic barrier function. We then challenged mice treated systemically with vancomycin in a model of dextran sulfate sodium (DSS)-induced colitis. We reasoned that this is the most suitable model of intestinal inflammation in this study, because DSS directly impairs the colonic mucus layer and severely impacts mice with an impaired mucus barrier^14,20^. We treated PBS- and vancomycin-treated mice with 4% DSS in drinking water, which results in only mild colitis in Swiss Webster mice. We found that vancomycin-treated mice lost more weight and showed more severe signs of disease that control mice (Figure 4D and E). Importantly, vancomycin-treated mice had strikingly larger colonic areas with ulcers than control mice (Figure 4F and G), indicating poor protection by the mucus layer against the DSS. Thus, vancomycin impairs mucus secretion from colonic goblet cells in a microbiota-independent manner which damages host protection against gut inflammation.

**Figure 4:**
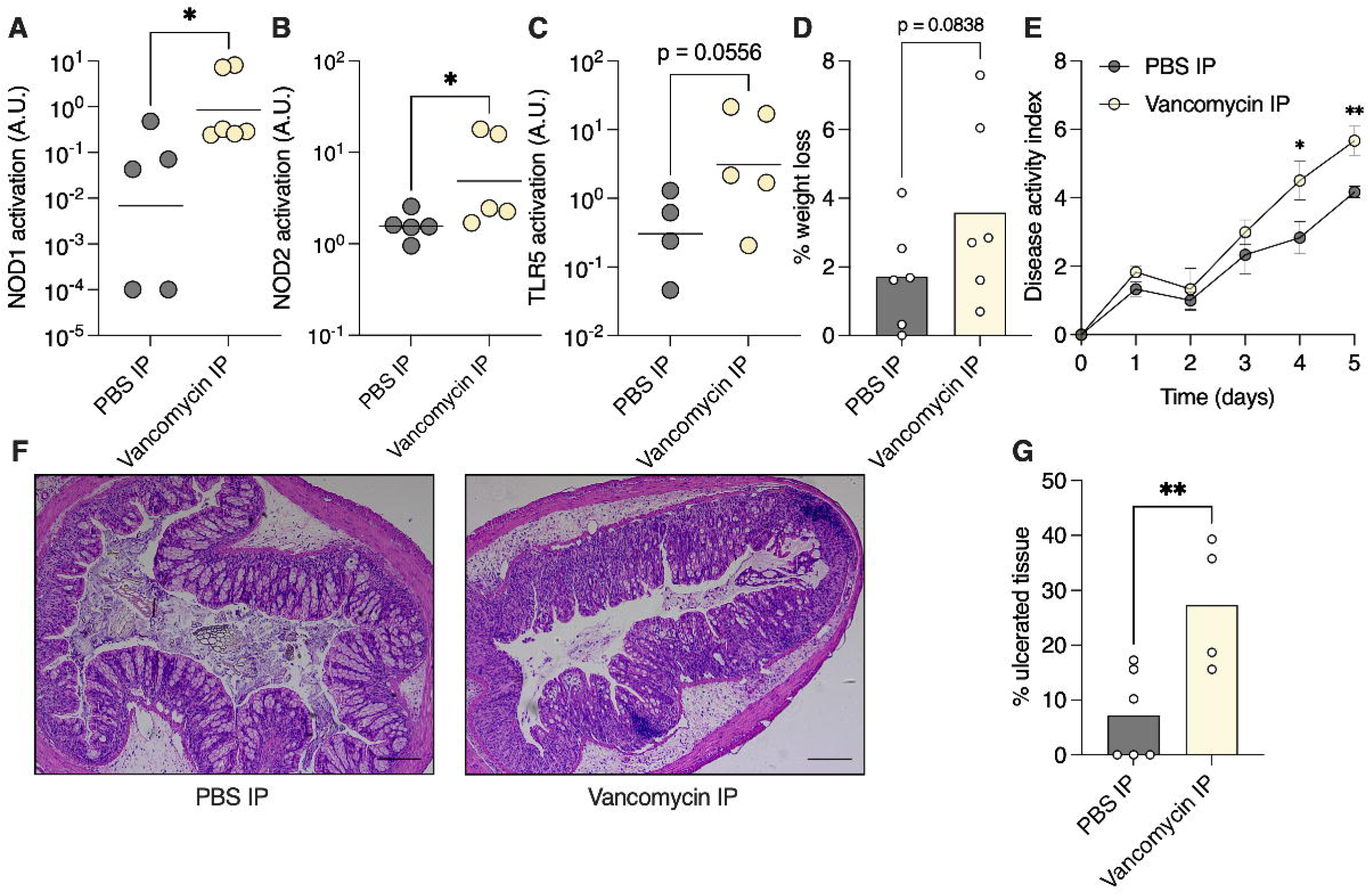
Systemic vancomycin treatment impairs colonic barrier function and aggravates colitis. (**A**-**C**) Detection of NOD1 (**A**), NOD2 (**B**) and TLR5 (**C**) agonists in mouse serum using reporter cell lines. (**D**-**G**) Mice were treated with 4% DSS for 5 days. (**D**) % weight change on day 5 of treatment compared to before treatment, (**E**) disease activity index ± SEM, (**F**) representative histological images of colonic tissue from DSS-treated mice and (**G**) quantification of percent of colonic area with ulceration. (**A**-**E** and **G**) Each dot represents a mouse. Scale bars, 20µm. (**A**-**E** and **G**) Student’s *t* test. *p<0.05, **p<0.01.

## Discussion

Antibiotics were hailed as miracle drugs, making once fatal infections seem mundane and casual. This success of antibiotics in improving human lives has led to their overuse in both medicine and agriculture. For a person living in the industrialized world, avoiding antibiotic exposure has become very difficult. The consequences of this overuse and overexposure has led to the rise of antibiotic-resistant pathogens, and, according to casual association, also development of a plethora of chronic diseases. Diseases such as diabetes, growth defects and chronic inflammations have been linked in multiple studies to antibiotic use^25–27^. As antibiotics target microbes directly, it is widely assumed that the effects of antibiotics on the gut microbiota are the main driver of these diseases^1,2^.

While antibiotic use affects the host by disrupting its microbiota, recent evidence suggests that antibiotics might act directly on host cells. By using germ-free mice, a few groups have shown that antibiotic drugs can activate an antiviral response^6^, affect tolerance to infection^5^ and influence host metabolism^4^ in a microbiota-independent manner by acting directly on host cells. How antibiotics exert this effect on host cells is not completely clear, yet it is thought they can interfere with mitochondrial function, DNA replication and various other cellular processes^3^. Even the use of antibiotics to protect cultured cells from infection, a staple of cellular and molecular biology, has come into question because of their effect on these cells^28^.

Another group of chronic diseases that are linked to antibiotic use are IBDs. Recent epidemiological studies have provided a strong link between antibiotic use and risk for development of IBD^9^, in a dose-dependent manner^10^. Here, we hypothesized that antibiotic use might lead to development of intestinal inflammation by affecting the colonic mucus barrier. The ability of the mucus barrier to provide separation between the host and its gut microbiota is crucial for maintaining gut homeostasis^12^. Indeed, breakdown of this barrier is observed both in animal models of IBD and in IBD patients^14,15^. As IBDs are characterized by loss of tolerance to the gut microbiota^29^, it is thought that impairment of the protective mucus barrier can drive these diseases. Without a proper mucus barrier, the microbes come in close contact with host tissues, triggering an immune response^13^.

We found that short-term oral antibiotic treatment was sufficient to impair the separation between host and microbiota in the colon (Figure 1). This phenomenon was true for all the antibiotics we tested. Using FMT and RNA sequencing followed by machine learning, we concluded that the effects of vancomycin on the mucus barrier could not be explained by antibiotic-induced changes to the microbiota alone (Figure 2). Instead, we found that vancomycin could impede mucus secretion in the colon, in a microbiota-independent manner, by inducing ER stress in colonic cells (Figure 3). Surprisingly, this effect of vancomycin on the ability of goblet cells to secrete mucus was immediate, as a few minutes following vancomycin infusion into the *ex vivo* chamber we observed a stark reduction in mucus secretion rates. Thus, we conclude that antibiotics have a deleterious effect on the mucus barrier, in part by acting directly on host cells. It is important to note that we do not rule out microbiota-dependent deleterious effects on the mucus barrier. But rather, that a direct effect on the host also exists. Indeed, a recent study has shown that transferring microbiota from humans with a history of antibiotic use to mice can cause mucus barrier defects in mice^30^.

Our study answers one question (does antibiotic-treatment impair the mucus barrier?), but raises two more questions. The first is: does antibiotic-treatment play a causative role in development of IBD? This question will be hard to answer in humans and will need to be tested in animal models of IBD. The major caveat of this approach is that animal models only partially reflect IBD pathology and development^31,32^. The second question is: how do antibiotics impair mucus production? Our experiments suggest that certain antibiotics induce ER stress in colonic cells, thus diminishing mucus production through a previously described mechanism^20,33^. We found it surprising that two distinct antibiotics from different classes, neomycin and vancomycin, both induced an ER stress response in the colon. As both drugs have distinct antimicrobial mechanisms, it is not clear why both would induce ER stress in host cells. It would be interesting to test whether antibiotic-treatment increases the risk for developing IBD in patients carrying predisposing mutations in autophagy-related genes, as autophagy is needed to relieve ER stress in goblet cells to allow proper mucus secretion^20^. Indeed, mutations in autophagy-related genes are associated with development of IBD^8,34^.

## Acknowledgment

This work was supported by the Azrieli Foundation Early Career Faculty Fellowship (to SB), the Israeli Science Foundation (ISF) (925/19 and 1851/19 to SB), the European Research Council (ERC) Starting Grant (GCMech 101039927 to SB) and Swedish Research Council grants 2018-02095 and 2021-06602 (to BOS).

## Materials and methods

### Ethics statement

All experiments in mice were conducted in compliance with the EU directive regarding the protection of animals used for experimental and other scientific purposes. Experiments were approved by the institutional animal care and use committee (IACUC) of the Bar-Ilan University (study ID #25-04-2019). Animal experiments performed at Umeå University, Sweden, were approved by the local animal ethical committee (Dnr A14-2019).

### Mice

C57BL6/J and Swiss Webster mice were bred and maintained under a 12 hour light/dark cycle and fed standard chow in either SPF or germ-free conditions at the Azrieli Faculty of Medicine of the Bar-Ilan University. For ex-vivo mucus measurements, C57BL6/J mice originally obtained from Charles River Laboratory Germany were bred in-house at Umeå university, Sweden, and maintained under a 12 hour light/dark cycle in individually ventilated cages in a pathogen-free environment at 22±1°C. Mice were fed a standard chow diet and had *ad libitum* access to food and water (#801730, Special Diet Services, UK).

### Antibiotic and TUDCA treatments

8 to 14-week-old mice were treated twice a day for 3 days with 2.5 mg of either ampicillin, metronidazole, neomycin or vancomycin dissolved in 100µl drinking water (for oral treatment) or PBS (for intraperitoneal injection). On the fourth day mice were euthanized according to IACUC guidelines and tissues and blood were harvested. For TUDCA treatment, mice were treated via intraperitoneal injection with 250 mg/kg TUDCA (Sigma T0266) dissolved in PBS twice daily in addition to vancomycin treatment.

### Fluorescent *in situ* hybridization (FISH), mucus thickness and bacterial distance from epithelium measurements

Mid-colon tissues containing a fecal pellet were excised from euthanized mice and immediately fixed in Carnoy’s fixative to preserve the mucus layer. Tissues were then processed for paraffin embedding using a standard automated protocol. 7µm-thick section were deparaffinized and FISH was conducted according to standard protocol^35^ with the following probes: Pan-bacterial probes EUB338I (GCT GCC TCC CGT AGG AGT), EUB338II (GCA GCC ACC CGT AGG TGT) and EUB338III (GCT GCC ACC CGT AGG TGT); Gammaproteobacteria probes GAM42a (GCC TTC CCA CAT CGT TT) and BET42a (GCC TTC CCA CTT CGT TT); Clostridia Clep866 (GGT GGA TWA CTT ATT GTG) and Erec482 (GCT TCT TAG TCA RGT ACC G). Slides were mounted with a DAPI-containing mount. Images were captured using an AxioImager M2 fluorescent microscope and distance between the host epithelium and bacteria quantified using Zeiss Zen Blue software. For mucus thickness measurements sections were stained with Alcian blue and mucus thickness was measured according to standard protocol^20^.

### Fecal microbiota transplant

Feces collected from vancomycin-treated mice housed at the SPF barrier facility were immediately transferred into an anaerobic chamber. Feces were vortexed in sterile PBS, debris allowed to settle by gravity, and supernatant transferred to new tubes. Sealed tubes were then removed from the anaerobic chamber and orally administered to germ-free Swiss Webster mice via gavage. Inoculated mice were then transferred to Tecniplast IsoCages to prevent outside contamination for 24 hours before tissue collection.

### RNA sequencing

RNA from frozen colonic tissues was extracted using Qiagen RNeasy Universal kit. Integrity of the isolated RNA was analyzed using the Agilent TS HS RNA Kit and TapeStation 4200 at the Genome Technology Center at the Azrieli Faculty of Medicine, Bar-Ilan University, and 1,000ng of total RNA was treated with the NEBNext poly (A) mRNA Magnetic Isolation Module (NEB, #E7490L). RNA sequencing libraries were produced by using the NEBNext Ultra II RNA Library Prep Kit for Illumina (NEB #E7770L). Quantification of the library was performed using a dsDNA HS Assay Kit and Qubit (Molecular Probes, Life Technologies) and qualification was done using the Agilent TS D1000 kit and TapeStation 4200, and 250nM of each library was pooled together and diluted to 4nM according to the NextSeq manufacturer’s instructions; 1.6pM was loaded onto the Flow Cell with 1% PhiX library control. Libraries were sequenced with the Illumina NextSeq 550 platform with single-end reads of 75 cycles according to the manufacturer’s instructions.

### Mapping RNA sequences for transcriptome analysis

The RNA sequencing data were analyzed using the RASflow pipeline^36^ mapping the reads to the transcriptome (Mus_musculus.GRCm39.cdna) using the hisat2 aligner^37^ and counting features using featureCounts^38^. For the genome and transcriptome datasets, any missing gene names (using mmusculus_gene_ensembl) are replaced with gene IDs.

We filtered out sparse genes, which we defined as those with over 50% zero values, to concentrate on genes with meaningful expression levels and to decrease computational complexity. We started with 35453 genes and removed 14981 sparse genes, leaving us with 20472 genes. Following this, we normalized the data with a scaling factor of 10^6^. The purpose of this normalization was to make the samples comparable by compensating for variations in library sizes. PCA plot was generated using RNAlysis^39^. Pathway analysis was performed using Enrichr^40–42^. The RNA sequencing data is available at NCBI GEO GSE260592.

The metadata, RASflow configuration scripts and relevant outputs including the normalized and unnormalized transcript and gene counts can be found in the github repository: https://github.com/AmirErez/Manuscript-Antibiotics_Damage_The_Colonic_Mucus/

### Four-way random forest classification

To accurately distinguish among the four groups: Van donor, Van recipient, PBS donor, and PBS recipient, we trained a random forest classifier. The classifier was trained using 20 samples, with 5 from each group, and tested on 8 samples, 2 from each group. The features were normalized to ensure each row’s sum equaled one million, standardizing the data scale for more effective learning. Using the RandomForestClassifier from scikit-learn, we configured each forest with 200 estimators (trees) and conducted 10,000 iterations.

The scripts for the classification can be found in the github repository: https://github.com/AmirErez/Manuscript-Antibiotics_Damage_The_Colonic_Mucus/

### Barrier function analysis

Blood was collected post-mortem via cardiac puncture and incubated at room temperature for 30 minutes in 1.5ml tubes to allow clotting. Samples were then centrifuged at 1,500g for 20 minutes at 4°C after which serum was collected to new tubes. 20µl of serum was added to wells containing InvivoGen HEK-Blue reporter cells, and luminal antigens were detected following manufacturer’s instructions.

### Barrier function analysis

Blood was collected post-mortem via cardiac puncture and incubated at room temperature for 30 minutes in 1.5ml tubes to allow clotting. Samples were then centrifuged at 1,500g for 20 minutes at 4°C after which serum was collected to new tubes. 20µl of serum was added to wells containing InvivoGen HEK-Blue reporter cells, and luminal antigens were detected following manufacturer’s instructions.

### Ex vivo mucus measurements and vancomycin treatment

Mucus growth rate in colonic tissue explant was measured as previously described^43^. Briefly, distal colon tissue from 8-12 week-old wild type mice was collected and washed with 4-5 ml of Kreb’s transport buffer to remove luminal content and unattached mucus. After muscle layer removal, the tissue was separated into 2 pieces. The tissues were mounted in horizontal perfusion chambers and maintained at 37°C. One piece of colon was incubated basolaterally with RPMI supplemented with 1.25 mg/ml vancomycin, while the other was mounted and incubated basolaterally with RPMI, as a control. The mucus was overlaid with 10 μm-sized beads to visualize the surface and the mounted tissues were then covered by Kreb’s mannitol buffer to maintain a moist environment. Mucus thickness was measured repeatedly with a micromanipulator-connected glass needle over 45 min, and the mucus growth rate (μm/min) was calculated as the change in mucus thickness per minute.

### Chemical-induced colitis model

Swiss Webster housed under SPF conditions mice were treated with 4% dextran sulfate sodium (DSS, colitis grade, 36,000–50,000 Da, MP) in drinking water. Fresh DSS was prepared daily. Disease activity index (DAI) was measured daily, based on weight loss, stool consistency and rectal bleeding as previously described^35^. Briefly, weight loss relative to initial weight, stool consistency (solid, loose or diarrhea) and rectal bleeding were each individually scored on a 0-4 scale and summed for each mouse at the indicated time points. For measurements of ulceration area, H&E-stained colonic sections from treated mice were visualized using a light microscope as above. Ulcerated surface area and healthy area were measured using Zeiss Zen Blue software. % ulcerated area was calculated as [ulcerated area]/[healthy area+ulcerated area].

## Supplementary Figures

**Figure S1:**
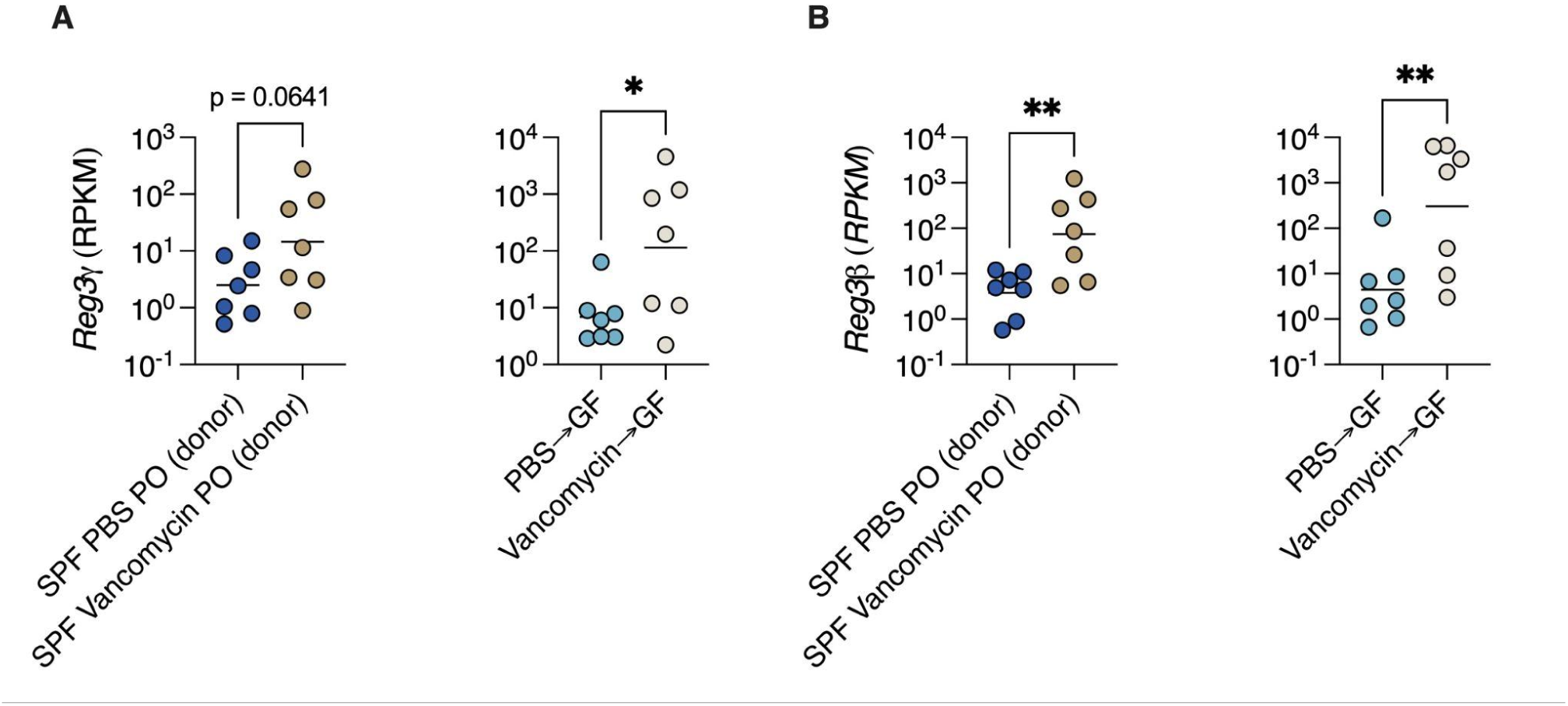
Antimicrobial genes are induced by vancomycin-altered microbiota. (**A**-**B**) Normalized reads of antimicrobial genes (**A**) *Reg3Ɣ* and (**B**) *Reg3*β in mice treated as indicated. Each dot represents a mouse. Student’s *t* test. *P<0.05, **P<0.01.

